# Inhibitory network predicts microstimulation-induced circuit changes in the awake mammalian cortex

**DOI:** 10.64898/2026.03.15.711307

**Authors:** Ming W. H. Fang, Maria C. Dadarlat, Yujiao Jennifer Sun

## Abstract

Intracortical microstimulation is a powerful tool to perturb neural circuits, yet how circuit changes post-stimulation remains poorly understood. Using two-photon imaging in the awake mouse visual cortex, we tracked genetically identified neurons before, during, and after microstimulation. Even after a short 15-min microstimulation, there is pronounced suppression in excitatory neurons, accompanied by increased activity in inhibitory neurons, particularly those not recruited during microstimulation. Our findings highlight inhibition as a key player in shaping stimulation-induced circuit changes: the magnitude of plasticity in excitatory neurons is not only dependent on the level of recruitment during stimulation, but also shaped by the recruitment level of neighboring inhibitory cells, whereas the inhibitory plasticity is best explained by pre-stimulation population coupling.

## Main

Electrical microstimulation has long been recognized for its high efficiency and spatiotemporal precision in driving neural activity, both in clinical and research applications. As the field marked the centenary of the first human intracortical microstimulation studies, which evoked artificial sensations (“phosphenes”) by delivering the electric current in patients’ visual cortices^1–3^, recent advances in electrode engineering and stimulating algorithms^4,5^ has renewed interest in intracortical microstimulation for neuroprosthetic and brain–machine interfaces^6,7^. A long-standing challenge, however, is that stimulation is delivered into a highly dynamic system: internal state and ongoing cortical activity strongly shape stimulation-evoked responses and the network activity^8–10^. Adding to this complexity, neurons in the cortex are heterogeneous, and microstimulation recruits a sparse, distributed population in a cell-type-specific manner^8,11,12^. Such selective recruitment can disproportionately engage specific excitatory and inhibitory populations, thereby shifting the excitation-inhibition balance that gates long-term plasticity^13–15^. Thus, designing reliable stimulation with predictive outcome requires accounting for the circuit’s dynamic and adaptive state and identifying the rules that link stimulation to both immediate recruitment and its after-effects. Here, we addressed these questions by tracking genetically identified excitatory and inhibitory neural populations at single-cell resolution before, during, and after microstimulation.

We performed two-photon calcium imaging in adult mice expressing the pan-neuronal calcium indicator GCaMP6s in layers 2/3 of the mouse primary visual cortex (V1), with inhibitory neurons labelled with red fluorescence Tomato in a GAD2-ires-cre;Ai14 background. Mice were awake and head-fixed on a floating ball, freely to run or stand still. Microstimulation was delivered through a single electrode, with trains of biphasic pulses at varying current amplitudes delivered in pseudo-random order (3–50 *μ*A; details in Methods). Imaging was acquired across three planes centered around the stimulating electrode, and neurons were tracked and matched across epochs within the same field of view. In total, we followed 1,155 excitatory neurons and 98 inhibitory neurons across four mice before, during, and after microstimulation (Fig. 1A).

**Figure 1:**
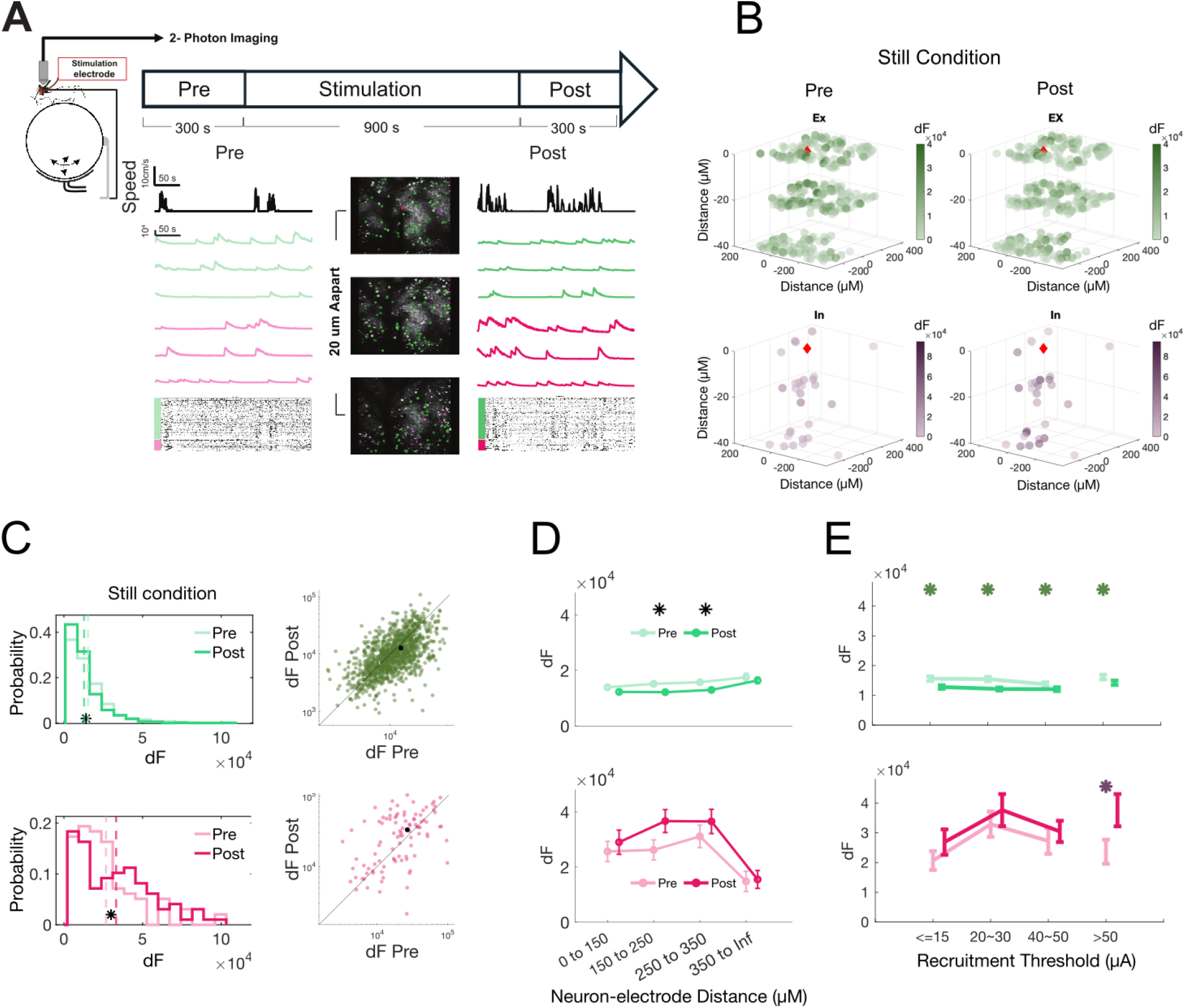
Two-photon imaging of excitatory and inhibitory neuronal populations before and after microstimulation. A) Experimental timeline and setup. Head-fixed, awake mice freely transitioned between locomotion and quiescence while three imaging planes (20 *μ*m apart) in layer 2/3 of V1 were imaged before stimulation (300 s), during stimulation (900 s), and after stimulation (300 s). A chronically implanted Pt/Ir electrode delivered pseudo-randomized current pulses (3–50 *μ*A) at 10 s intervals. Example calcium traces (*dF*) and treadmill speed are displayed. B) Example field of view from one mouse showing excitatory (green) and inhibitory (red) neurons across three imaging planes before and after stimulation (still condition shown). The position of the electrode is denoted by the red diamond. C) Distribution of mean still-state *dF* activity before and after stimulation for excitatory (top) and inhibitory (bottom) neurons pooled across mice. Left: staircase histograms. Right: pre vs post scatter plots. Excitatory neurons show significant post-stimulation suppression, whereas inhibitory neurons show increased activity. Statistical comparisons were performed using paired tests across neurons; significance indicated by black stars (\**p <* 0.05). D) Mean still-state *dF* as a function of neuron–electrode distance (*μ*m). Neurons were grouped into distance bins (0–150, 150–250, 250–350, *>* 350 *μ*m). Excitatory neurons were significantly suppressed at intermediate distances (150 to 350 *μ*m, *p*s *<* 0.01). E) Mean still-state activity before and after stimulation binned by recruitment threshold (3–15, 20–30, 40–50, *>* 50 *μ*A). Excitatory neurons showed consistent suppression across thresholds. Inhibitory neurons showed increased activity in the *>* 50 *μ*A (never recruited) bin. Stars indicate significant pre–post differences (\**p <* 0.05). All error bars denote SEM.

### Cell-type-specific modulation of activity after micro-stimulation

We compared the neural activity before and after microstimulation during animals’ quiescent epochs (speed *<* 1 cm/s), to isolate stimulation-driven changes from modulation of cortical activity by locomotion^16,17^. Microstimulation induced robust, cell-type-specific changes in neural activity (Fig. 1C); across the population, the mean activity of excitatory neurons was suppressed post-stimulation, whereas inhibitory neurons were more active. Because microstimulation recruits neuronal populations with distinct spatial profiles^8,12^, we first grouped neurons based on their neuron-electrode distance and compared their activity before and after stimulation: excitatory neurons at intermediate distances (150–350 *μ*m) were most significantly suppressed, while inhibitory neurons were weakly, but not-significantly more active within 350 *μ*m of the stimulating electrode (Fig. 1B, D).

We further grouped neurons based on their recruitment threshold: because the stimulation protocol spanned nine amplitudes (3–50 *μ*A), neurons with higher recruitment thresholds would be recruited less often, or not at all, during the stimulation phase. Excitatory neurons, regardless of its recruitment level, including non-recruited cells (*>*50 *μ*A), were consistently suppressed. In contrast, recruited inhibitory neurons exhibited only modest changes while non-recruited inhibitory neurons had significantly enhanced activity (Fig. 1E), suggesting their potential contribution to the circuit changes.

### Local inhibitory recruitment pattern governs excitatory plasticity

To quantify the level of activity changes, we computed an electrical modulation index (EMI) for each neuron, defined as the normalized change in average activity during quiescence before and after stimulation (see Method). In excitatory neurons, EMI was predominantly negative (Fig. S1A) and showed a modest dependence on recruitment threshold, with more strongly recruited neurons tending to exhibit greater post-stimulation suppression (more negative EMI; Fig. 2A). This relationship suggested that stimulation history contributes to the post-stimulation changes at the level of individual neurons.

**Figure 2:**
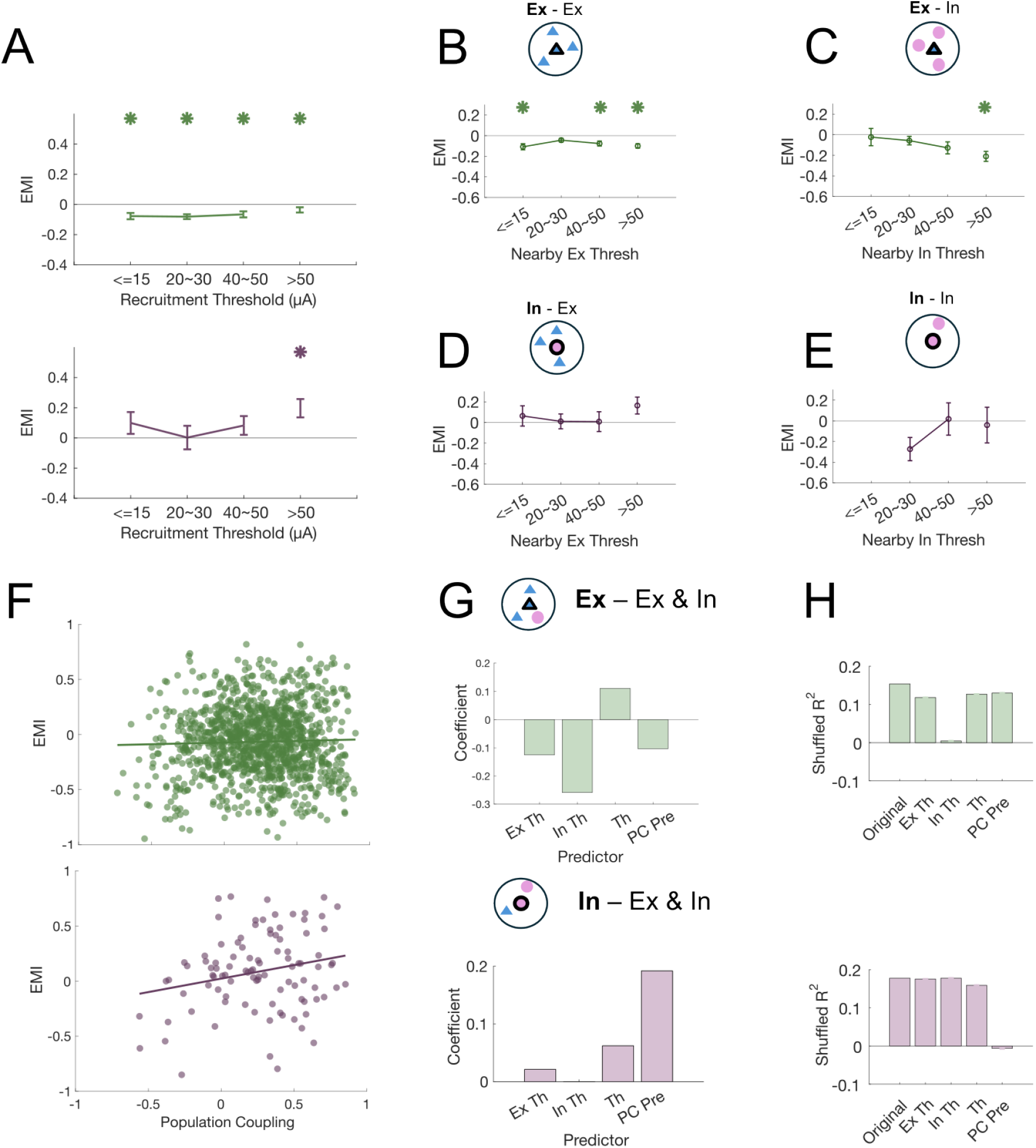
Cell-type-specific electrical modulation correlates with nearby cells’ activation pattern and their own network connectivity. A) Electrical modulation index (EMI = (*dF*_post_ *− dF*_pre_)*/*(*dF*_post_ + *dF*_pre_)) during still state for excitatory (top) and inhibitory (bottom) neurons binned as a function of recruitment thresholds. Excitatory neurons show suppression across all threshold bins. Inhibitory enhancement is most pronounced in the *>* 50 *μ*A (never recruited) group. All one-sample *t* tests *p*s *<* 0.01. B–E) Electrical modulation index of cells as a function of nearby (*<* 25 *μ*m) cells’ recruitment thresholds, separated by center-neighbor identity: B) Ex–Ex, C) Ex–IN, D) IN–Ex, E) IN–IN. Excitatory neurons surrounded by inhibitory neighbors (Ex–IN) show stronger suppression when nearby inhibitory neurons were weakly recruited (higher recruitment threshold), all *p*s *<* 0.01. Same-type neighborhood configurations show minimal predictive structure. Error bars denote SEM. F) Relationship between pre-stimulation population coupling and EMI for excitatory (top, slope = 0.03, *R*^2^ = 0, correlation coefficient = 0.03, *p >* 0.05) and inhibitory (bottom, slope = 0.24, *R*^2^ = 0.05, correlation coefficient = 0.23, *p <* 0.05) neurons. Each dot represents one neuron. EMI is significantly correlated with pre-stimulation coupling only in inhibitory neurons. G) Multi-variate regression model predicting EMI from four predictors: nearby excitatory recruitment threshold (Ex Th), nearby inhibitory recruitment threshold (In Th), own recruitment threshold (Th), and pre-stimulation population coupling (PCPre). Bars show standardized regression coefficients. H) Change in model performance (Δ*R*^2^) following shuffling of each predictor. For excitatory neurons, nearby inhibitory threshold contributes most strongly to predictive power. For inhibitory neurons, PCPre dominates. Error bars rep1re2sent SEM.

Because individual neurons receive dense local synaptic input, we next tested whether EMI is also influenced by the nearby neurons. Excitatory EMI was strongly influenced by local excitatory–inhibitory context: excitatory neurons surrounded by less-recruited inhibitory neighbors (i.e., inhibitory neurons with higher recruitment thresholds) were more strongly suppressed. No clear relationship was observed with nearby excitatory neighbors, despite their tendency to share similar recruitment levels (Fig. 2B–C, Fig. S1B–D). In contrast, inhibitory EMIs was relatively independent of neighboring recruitment level (Fig. 2D–E, Fig. S1E–F).

### Inhibitory plasticity reflects intrinsic network coupling

Given that microstimulation response depend on pre-exising network activity, we further tested whether post-stimulation changes depend on properties that are present even before microstimulation. To address this, we calculated pre-stimulation population coupling (Fig. S2), a metric that reflects the amount of local synaptic input in cortical networks, by correlating a neuron’s activity with the overall population activity.^18^ While population coupling did not predict EMI for excitatory neurons, we found a significant positive relationship between population coupling and inhibitory neuron EMI (Fig. 2F), especially in the stimulation-recruited population (Fig. S3).

To assess the relative contributions of candidate determinants for post-stimulation changes, we fitted a multivariate regression model to predict each neuron’s EMI as a function of its recruitment threshold (“Th”), the recruitment level of nearby excitatory neighbors (“Ex Th”), the recruitment level of nearby inhibitory neighbors (“In Th”), and the neuron’s pre-stimulation population coupling (“PC Pre”; Fig. 2G). For excitatory neurons, nearby inhibitory neighbor recruitment was the strongest predictor of post-stimulation activity changes, as indicated by largest regression weight and a marked reduction in model performance (*R*^2^) when this predictor was shuffled (Fig. 2H). In contrast, for inhibitory neurons, population coupling had the most predictive power, whereas recruitment patterns contributed minimally. Together, these results indicate that excitatory plasticity is governed by local inhibition during stimulation, whereas inhibitory plasticity is determined by intrinsic network connectivity.

### Microstimulation reorganizes network connectivity

Next, we asked whether microstimulation alters population coupling, a measure of local network connectivity, and whether such changes depend on recruitment during stimulation. Excitatory neurons showed a significant increase in population coupling following stimulation (Fig. 3A). This increase was most prominent in strongly recruited neurons (recruitment threshold *≤*15 *μ*A and 20–30 *μ*A), although pre-stimulation coupling was broadly similar across recruitment thresholds (Fig. 3B). In contrast, inhibitory neuron coupling was unchanged after stimulation (Fig. 3A,B). Notably, inhibitory recruitment tended to depend on initial population coupling: neurons with high pre-stimulation coupling were strongly recruited, whereas weakly coupled neurons were also only weakly recruited (40–50 *μ*A) or not recruited (*>* 50 *μ*A; Fig. 3B; see also Fig. S2B). These effects were not explained by neuron-electrode distance, as coupling did not vary systematically with spatial proximity to the electrode (Fig. S2C). Together, these results indicate that strongly recruited excitatory neurons during stimulation undergo greater enhancement in coupling post-stimulation, whereas inhibitory recruitment may depend in part on pre-existing network coupling.

**Figure 3:**
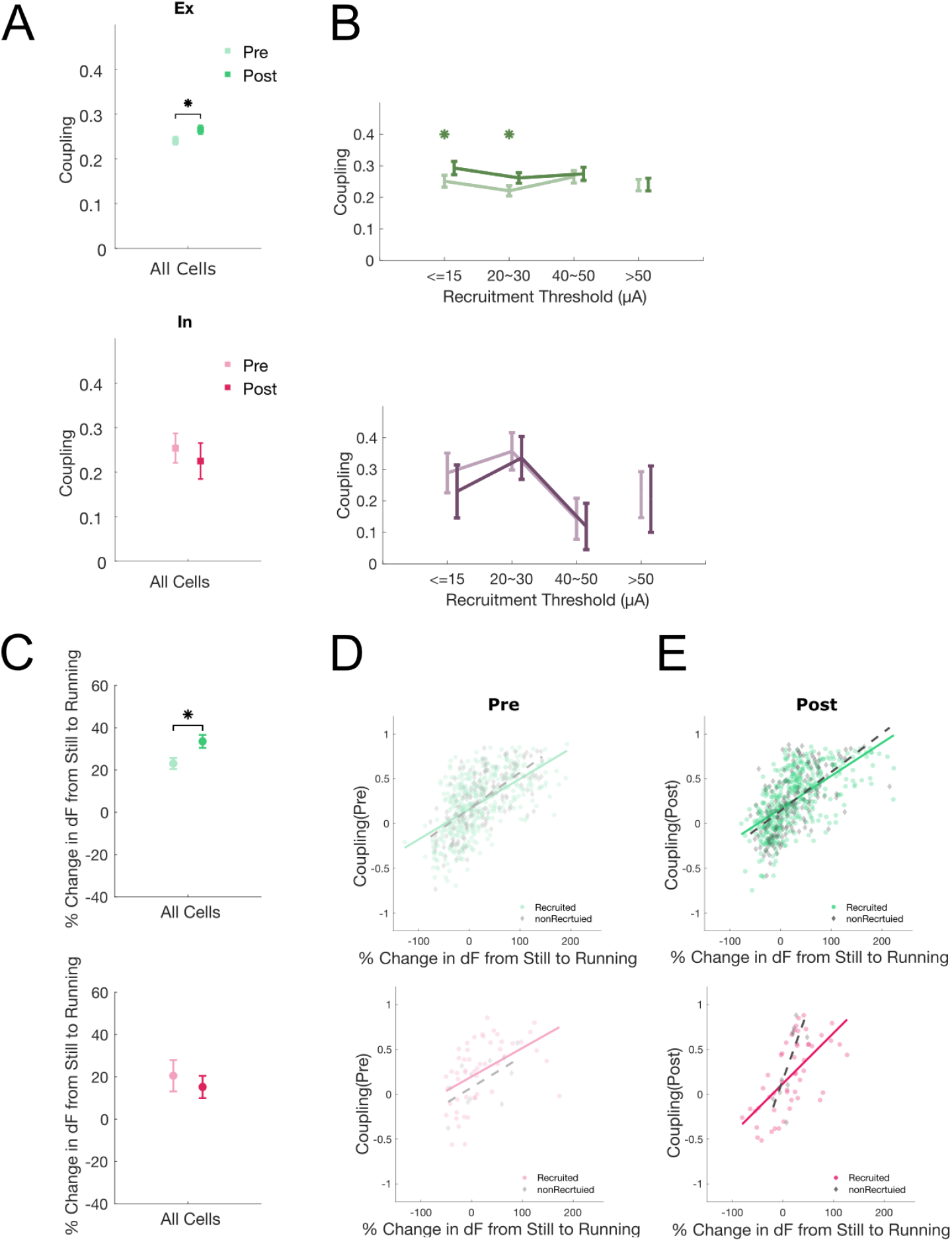
Microstimulation reorganizes network connectivity. A) Microstimulation increased coupling of excitatory neurons (top) within the network (*p <* 0.05). However, inhibitory neurons’ connectivity remained stable before and after stimulation. B) Population coupling of different recruitment threshold bins before and after stimulation. For excitatory cells, coupling remained similar across threshold bins both before and after stimulation (Kruskal–Wallis test, all *p*s *>* 0.05). However, strongly recruited cells (e.g., *≤*15, and 20–30 *μ*A) significantly increased their coupling after stimulation (paired *t* tests, *p*s *<* 0.05). On the contrary, for inhibitory neurons, there is a trend (Kruskal–Wallis test, *p* = 0.09) that more recruited neurons tend to have higher baseline coupling (i.e., “pre” condition). But, we found no significant difference between their pre- and post-stimulation coupling. C) Percentage change in activity (*dF*) from still to running. Microstimulation induced stronger behavioral modulation on excitatory neurons’ activity (paired *t* test, *p <* 0.01), whereas behavioral modulation on inhibitory neurons’ activity remained unchanged. D–E) Relationship between behavioral modulation (percentage change in *dF* from still to running) and population coupling before (D) and after (E) stimulation. Recruited (colored) and non-recruited (grey) neurons are shown separately. Linear fits are overlaid (solid line: recruited, dashed line: non-recruited). Slopes of linear fits remained the same pre- and post-stimulation for excitatory neurons (recruited: pre = 0.34, post = 0.37, non-recruited: pre = 0.41, post = 0.43). However, non-recruited inhibitory neurons show a marked post-stimulation change in slope (pre = 0.35, post = 1.58), and a much smaller increase for the recruit1e3d group (pre = 0.32, post = 0.56), where a steeper slope reflected reduced behavioral sensitivity relative to coupling. Error bars represent SEM.

Finally, we examined how the change of population coupling impacts the cortical modulation by behavioral state. Population coupling has been shown to capture not only local circuit connectivity but also sensitivity to diffuse, nonspecific inputs outside of the sensory cortices.^18^ We therefore quantified the change in neural activity when an animal transitions from stillness to running and to test if the level of population coupling correlates with level of behavioral state-modulation. Indeed, we found the increased activity during locomotion initiation was highly correlated with population coupling in both excitatory and inhibitory populations.^17,19^ After stimulation, excitatory neurons were more sensitive to behavioral state modulation (Fig. 3C), but the relationship between coupling and state modulation remained consistent(Fig. 3D-E). In contrast, the association between population coupling and state modulation was markedly weakened for inhibitory cells, particularly in the non-recruited population (Fig. 3D-E and Fig. S4). More broadly, microstimulation strengthens local coupling in excitatory neurons while altering how inhibitory neurons, particularly non-recruited ones, relate behavioral-state signals to network activity.

## Discussion

Building on previous observation of strong inhibitory recruitment during stimulation,^8,20,21^ we found pronounced, cell-type specific changes following stimulation: broad suppression in the excitatory neurons was accompanied by enhanced activity in inhibitory neurons, especially the ones not activated during stimulation. These changes did not scale systematically with neuron-electrode distances, supporting circuit-level adaptation rather than passive current spread.^12,22^ Our results show that the after-effects of microstimulation depend not only on neurons directly recruited during stimulation, but also on inhibitory-network properties that are present before stimulation begins.

This perspective extends prior works studying microstimulation-induced plasticity^23,24^ by highlighting the contribution of inhibitory networks: excitatory plasticity was shaped by local inhibitory recruitment during stimulation, whereas inhibitory plasticity was better predicted by pre-stimulation population coupling, which remains relatively stable post-stimulation. Our results support a growing consensus that inhibitory neurons are well positioned to stabilize cortical activity and gate plasticity in response to external perturbation.^13–15,18,25^

Overall, this work suggests that the outcome of a given microstimulation protocol is shaped not only by stimulation parameters, but also by the intrinsic network properties present before stimulation. Predicting stimulation efficacy will therefore require accounting for circuit state, particularly the organization of inhibitory populations, in addition to the choice of stimulation parameters. More broadly, defining the inhibitory circuit features that govern post-stimulation changes may help guide strategies to better harness, or mitigate, cortical reorganization in brain–machine interfaces and therapeutic stimulation protocols.

## Methods

### Animals

All procedures were approved by the Institutional Animal Care and Use Committee at the University of California, San Francisco and conformed to National Institutes of Health guidelines. Surgical preparation and chronic imaging protocols followed previously established procedures^8^ and are summarized briefly below.

Inhibitory neurons were genetically labeled by crossing *Gad2-IRES-Cre* mice with the Ai14 reporter line (Jackson Laboratory strains 010802 and 007914;^26,27^). Experiments were performed in four adult mice (3–6 months old; both sexes). Each animal underwent sequential surgical procedures for implantation of a titanium headplate, a chronic cranial window, and a platinum–iridium stimulating electrode.

Prior to electrode implantation, mice received virual injections of GCaMP6s (Addgene 100843-AAV1;^28^) into primary visual cortex (V1) at three sites and two depths (150 and 300 *μ*m) per site to label neuronal populations with a genetically encoded calcium indicator. Surgeries were performed under ketamine/xylazine anesthesia (100 mg/kg and 5 mg/kg, intraperitoneal).

### Surgical procedures

Headplate implantation and chronic window preparation followed previously described methods^8,16,29^. Briefly, a craniotomy was performed over V1, and a Pt/Ir microelectrode (0.1 MΩ; Microprobes for Life Sciences) was implanted at the edge of the cranial window. A shallow skull ramp was created anterior to the craniotomy to allow electrode insertion while preserving stable window placement. The electrode and window were secured to permit long-term imaging and stimulation in awake animals.

### Electrical stimulation

Electrical stimulation protocols were identical to those described previously^8^. During each stimulation session, constant-current stimuli were delivered once every 10 s at 9 different amplitudes (3, 5, 10, 15, 20, 25, 30, 40, or 50 *μ*A) pseudo-randomly, with ten repetitions per amplitude (90 stimuli total). Stimuli consisted of a train of 25 biphasic cathode-leading pulses delivered at 250 Hz for a total duration of 100 ms, with each pulse comprised of a 200 *μ*s cathodal phase, 200 *μ*s interphase interval, and 200 *μ*s anodal phase.

Charge density remained below established safety limits for cortical stimulation based on electrode surface area and pulse parameters. In contrast to prior work focusing on stimulation-evoked responses, the present study analyzed persistent changes in spontaneous activity measured before and after stimulation.

### In-vivo two-photon calcium imaging

Calcium imaging was performed using a resonant-scanning two-photon microscope (Neuro-labware) controlled by Scanbox software. GCaMP6s was excited at 920 nm using a Coherent Chameleon Ultra II laser. Emitted fluorescence was separated using 525/70 nm and 610/75 nm emission filters and detected with photomultiplier tubes (Hamamatsu R3896).

Recordings were obtained from awake, head-fixed mice free to run on an air-supported spherical treadmill in a darkened enclosure^30^. Imaging was performed using a 16 *×*water-immersion objective (0.8 NA; Nikon) at depths of 100–300 *μ*m below the pia, corresponding to layer 2/3 of V1.

Neural responses were imaged in three planes spaced 20 *μ*m apart in depth, covering an area of approximately 1136 *μ*m by 1083 *μ*m. Each plane was imaged at 5.17 Hz (15.5 Hz divided by three imaging planes). All neural recordings were made in a dark room and the animals were isolated from the experimenter in an enclosure formed by black laser safety fabric. Mice underwent at least 5 days of training to acclimate to the treadmill and headfixation prior to recordings. Running speed was recorded using a synchronized camera (Dalsa Genie M1280) tracking treadmill motion.

### Image acquisition and processing

Two-photon imaging data were processed using the Suite2p pipeline for motion correction, ROI detection, and neuropil correction^31^. The data were further curated through manual cell detection, differentiating between cell and non-cell ROIs using the Suite2p GUI. Each ROI identified by Suite2p was manually checked to determine whether Suite2p’s classification of cell or non-cell was correct. The classifier was trained by considering the calcium traces, shape, compactness, aspect ratio and position of the ROIs. ROIs were considered as cells if: 1) they had calcium traces considerably different from the surrounding neuropil activity, and 2) a round, compact shape of a size large enough to represent a cell body rather than the cross section of a dendrite. We kept all ROIs that were assigned by Suite2p a probability of at least 0.5 of being a neuron. All subsequent analyses were conducted in MATLAB using custom code.

Genetically labeled inhibitory neurons were identified from ROIs in the red channel using a modified implementation of the Suite2p pipeline. Residual contamination from green-to-red channel was corrected using an estimation based on the minimum red-channel value at each green-channel intensity. ROIs were classified as inhibitory if the corrected red fluorescence exceeded the local population mean by more than two standard deviations. All downstream analyses were performed using custom MATLAB and Python scripts.

### Neural data analyses

Neuronal activity was quantified using neuropil-corrected calcium fluorescence (*dF* ; Suite2p *F*_corr_), and analyses focused on spontaneous activity during quiescent behavioral epochs (speed *<* 1 cm/s) unless otherwise noted. For spatial distance analysis, we found the position of the electrode tip by visually inspecting the max projection image across the aligned imaging movie data. Within each imaging plane, a neuron’s position was said to be at the mean of the *x* and *y* pixels assigned to that ROI, which were then converted into *μ*m. We compute the neuron–electrode distance and neuron–neuron distance for each neuron as Euclidean distance:

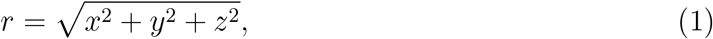

where *x* and *y* are the mean position of a neuron within the imaging plane (in mm) and *z* is the distance between the plane (0, 20, or 40 *μ*m).

Stimulation-induced plasticity was quantified using the electrical modulation index (EMI), defined as

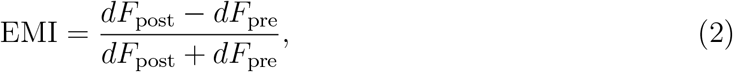

where *dF*_pre_ and *dF*_post_ denote mean *dF* during preand post-stimulation spontaneous still periods, respectively.

Recruitment status was determined by the lowest amplitude during microstimulation. Non-recruited neurons were never activated during the stimulation and assigned a threshold value of 100 *μ*A. Neuron–electrode distance was evaluated after binning into predefined ranges (0–150, 150–250, 250–350, *>* 350 *μ*m). Recruitment thresholds were grouped according to predefined stimulation bins (3–15, 20–30, 40–50, *>* 50 *μ*A).

Global network connectivity was quantified using population coupling, defined as the Pearson correlation between a neuron’s *dF* trace and the mean population trace excluding that neuron, computed separately for pre- and post-stimulation spontaneous periods. The coupling–plasticity relationship was assessed by correlating pre-stimulation coupling with EMI separately for each cell type. Multivariate regression models with L2 regularization (i.e., “Ridge” regression) included nearby excitatory threshold, nearby inhibitory threshold, own recruitment threshold, and pre-stimulation coupling to evaluate relative predictor contributions to EMI. We computed the shuffled *R*^2^ separately by shuffling one predictor (1000 repetitions) at a time.

Neighborhood similarity is determined by recruitment threshold and pre-stimulation population coupling. For threshold similarity, we derived:

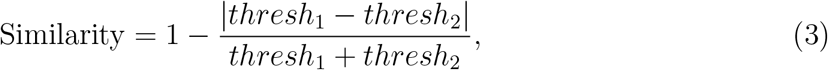

where *thresh*_1_ and *thresh*_2_ are the recruitment thresholds for any pair of neurons from the same imaging plane. For population coupling, we defined:

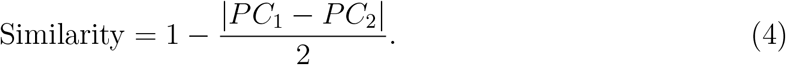

Similarity indices were sorted based on pairwise distance into predefined bins from 0 to 300 *μ*m at a step size of 25 *μ*m.

Behavioral state modulation was quantified as percentage of mean activity change from still to running, defined as

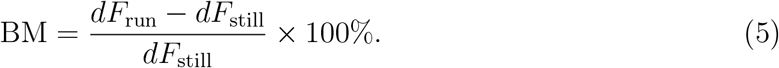

Microstimulation-induced changes in behavioral modulation were assessed using BM_post_*−* BM_pre_. Relationships between locomotion modulation and coupling were evaluated using ordinary least squares linear regression analyses performed separately by cell type and recruitment group.

### Statistics and reproducibility

Statistical analysis was performed using MATLAB (MathWorks). Group comparisons were performed using paired *t* tests and Kruskal–Wallis tests, unless otherwise specified. A Bonferroni correction was used to adjust for multiple comparisons. The alpha was set to 0.05 for all tests. Correlation analyses used Pearson correlation. Multivariate regression models were implemented using ridge regression. All statistical tests were two-sided. Error bars represent standard error of the mean. No statistical methods were used to predetermine sample sizes but our sample sizes are similar to those reported in previous publications in the field.

## Data and code availability

Data and custom analysis code will be made publicly available upon acceptance. Until then, they are available from the corresponding author upon reasonable request.

## Acknowledgement

We thank Professor Michael P. Stryker (UCSF) for support and access to laboratory resources during data acquisition. This work was supported by the IoO ECR Glaucoma Research fund (M.W.H.F), UCSF Program in Breakthrough Biomedical Research (M.C.D.), NSF HDR grant 2117997 (M.C.D), NIH grant 5K99EY029002 (Y.J.S.), and the Moorfields Eye Charity Springboard Award GR001264 (Y.J.S.).

## Author contribution

M.C.D., and Y.J.S. designed the study and performed the experiments. M.W.H.F., and Y.J.S. analyzed the data and wrote the manuscript. M.C.D., and Y.J.S reviewed the manuscript.

---

**Figure S1:**
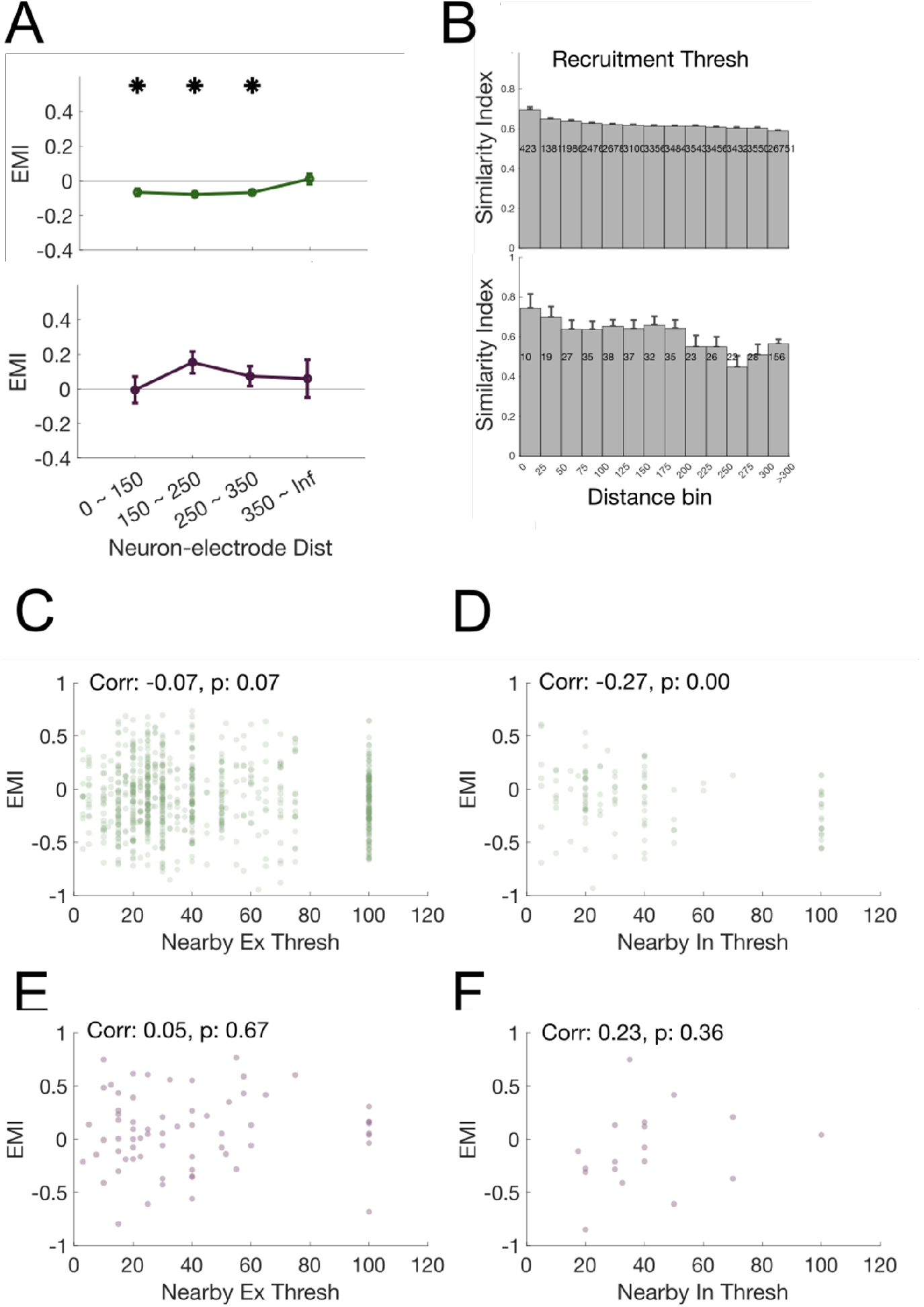
A) Electrical modulation index as a function of neuron–electrode distances (\**p <* 0.01). Error bars represent SEM. B) Neighbors’ similarity index using recruitment threshold plotted as a function of neuron–neuron distance. C–F) Electrical modulation index of cells as a function of nearby (*<* 25 *μ*m) cells’ recruitment thresholds, separated by center-neighbor identity: C) Ex–Ex, D) Ex–IN, E) IN–Ex, F) IN–IN. Excitatory neurons surrounded by inhibitory neighbors (D, Ex–IN) showed a significant negative correlation (*r* =*−* 0.27, *p <* 0.01) with the recruitment threshold of nearby inhibitory neurons. Same-type neighborhood configurations show minimal predictive structure.

**Figure S2:**
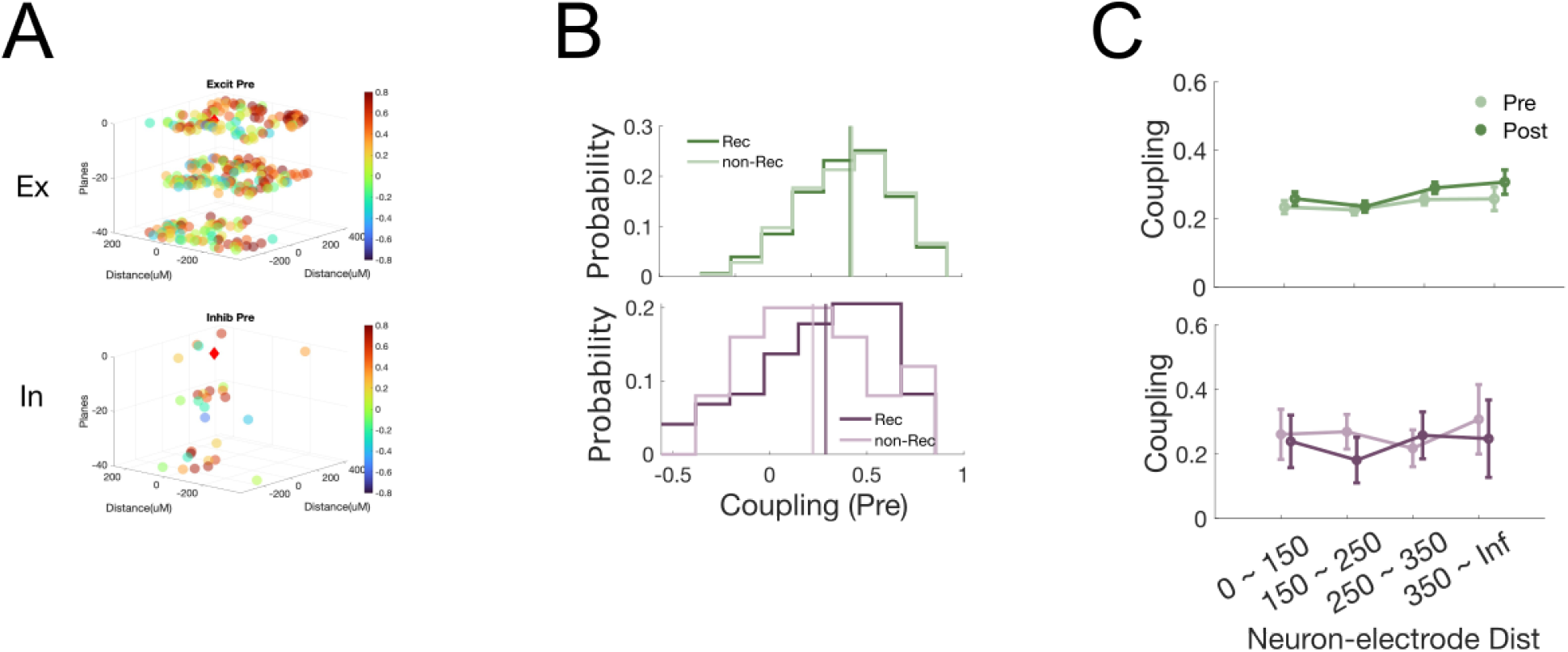
A) Example mouse’s spatial maps of population coupling across imaging planes for excitatory and inhibitory neurons. B) Distribution of pre-stimulation population coupling for recruited vs non-recruited neurons. C) Population coupling as a function of neuron–electrode distance (all neurons pooled). There is no significance across bins for pre and post for both cell types (Kruskal–Wallis test, *p*s *>* .05).

**Figure S3:**
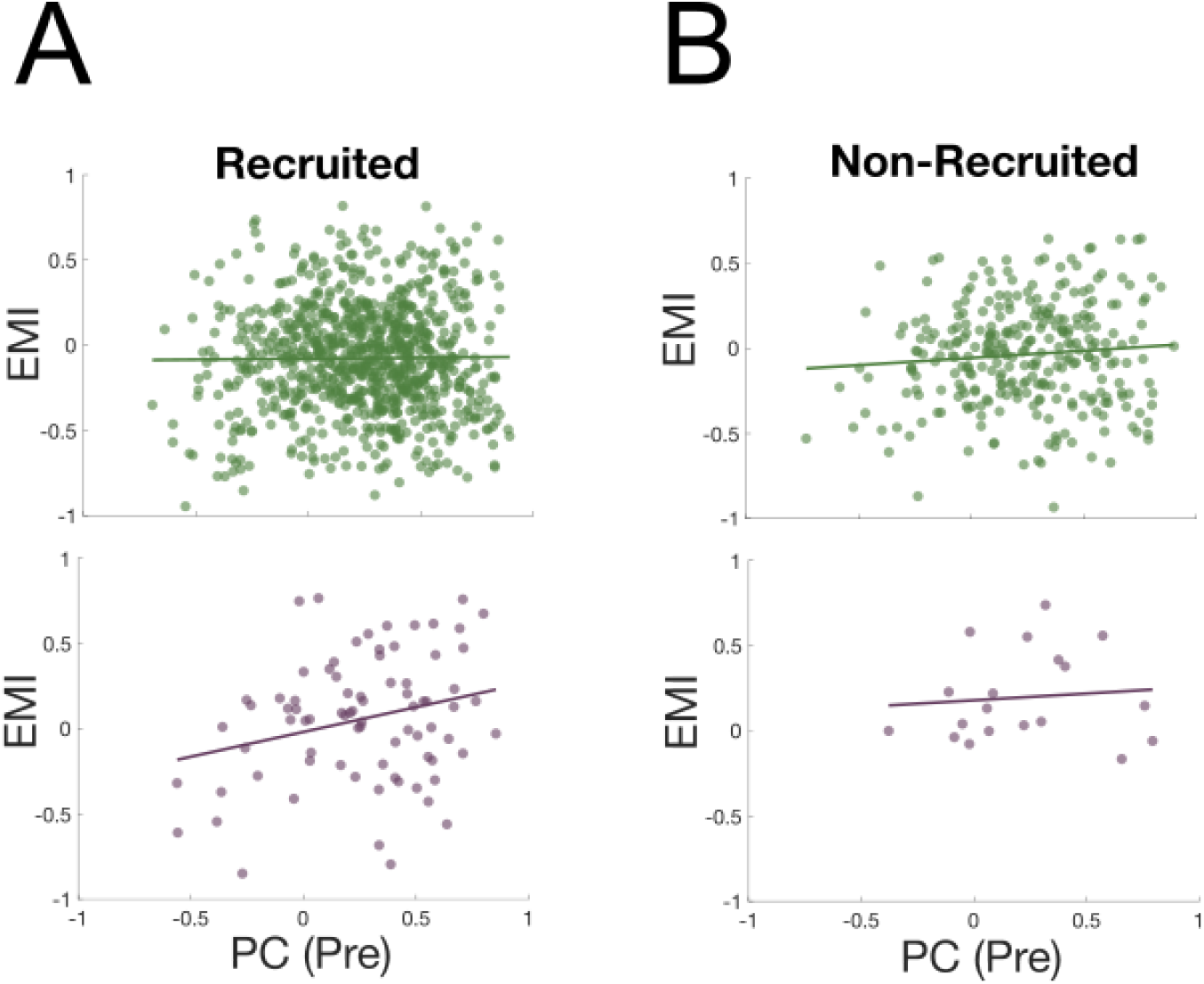
A–B) Relationship between pre-stimulation coupling and EMI separated by recruitment group for recruited (A) and non-recruited (B) cells. Recruited inhibitory cells governed the plasticity-coupling link (A, bottom).

**Figure S4:**
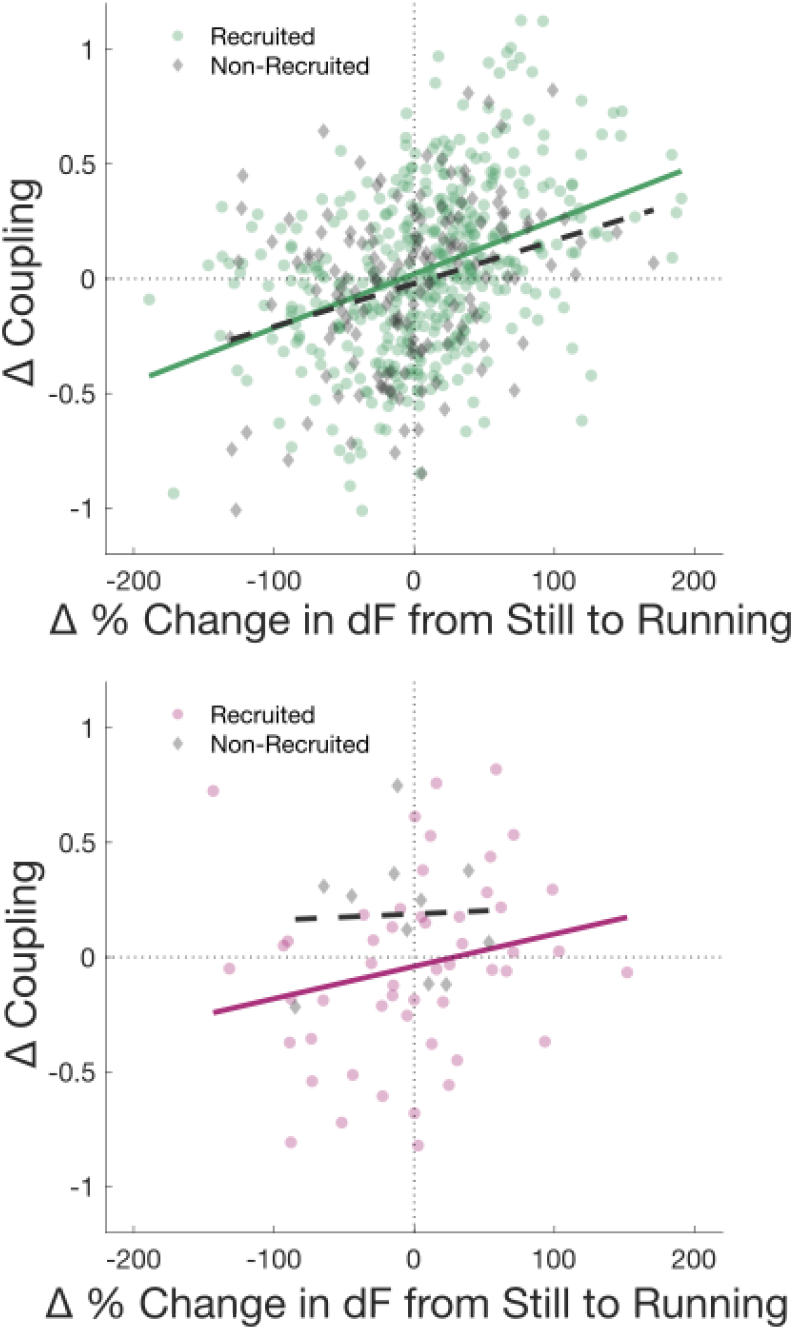
Δ PC plotted as a function of Δ % change in *dF* from still to running. For excitatory cells, the changes in coupling (post *−* pre) strongly correlated with Δ % change in *dF* from still to running for both recruited (linear fit slope = 0.24, correlation coefficient = 0.39, *p <* 0.01) and non-recruited (linear fit slope = 0.19, correlation coefficient = 0.3, *p <* 0.01) group. For inhibitory cells, there is a difference between recruitment status. The relationship is much reduced for non-recruited cells (linear fit slope = 0.027, correlation coefficient = 0.04) than the recruited group (linear fit slope = 0.14, correlation coefficient = 0.22).

